# Human transcription factor combinations mapped by footprinting with deaminase

**DOI:** 10.1101/2024.06.14.599019

**Authors:** Runsheng He, Wenyang Dong, Wenping Ma, Zhi Wang, Long Gao, Chen Xie, Dubai Li, Ke Shen, Fanchong Jian, Jiankun Zhang, Yuan Yuan, Xinyao Wang, Yuxuan Pang, Zhen Zhang, Yinghui Zheng, Shuang Liu, Cheng Luo, Xiaoran Chai, Jun Ren, Zhanxing Zhu, Xiaoliang Sunney Xie

## Abstract

An individual’s somatic cells have the same genome but exhibit cell-type-specific transcriptome regulated by a combination of transcription factors (TFs) for each gene. Mapping of TF sites on the human genome is critically important for understanding functional genomics. Here we report a novel technique to measure human TFs’ binding sites genome-wide with single-base resolution by footprinting with deaminase (FOODIE). Single-molecule sequencing reads from thousands of cells after *in situ* deamination yielded site-specific TF binding fractions and the cooperativity among adjacent TFs. In a human lymphoblastoid cell line, we found that genes in a correlated gene module (CGM) share TF(s) in their *cis*-regulatory elements to participate a particular biological function. Finally, single-cell resolved experiments (scFOODIE) allow cell-type-specific TF footprinting in heterogeneous brain tissues.

## Introduction

Every somatic cell in the human body has essentially the same genome but carries out a distinct function in a specific tissue with different gene expression. Transcription factors (TFs) bind to *cis*-regulatory elements, including promoters and enhancers of genes on the cell’s genome, to control gene expression and regulation, cell differentiation, and development, in response to environmental signals through regulatory networks (*1*). Our knowledge about gene expression and regulation first came from the study of bacteria, where one TF, acting as a key to turn on and off an operon, contains multiple protein-coding genes that carry out a particular biological function (*2, 3*). Such TF activities have been directly monitored and quantitatively described on a single-cell and single-molecule basis (*4, 5*).

In contrast, in humans, each gene encodes only one protein (and its isoformers). How are different proteins participating a complexed biological function, such as protein synthesis or glucose metabolism? TFs in eukaryotes, ever since their first isolation (*6*), seem to operate differently from bacteriain that each binds to many sites on the genome with lower binding affinities (*7*), and gene expression and regulation of a particular geneThe fact that several TFs, acting together as a set of keys, control a protein-coding gene makes the total number of ∼1600 TFs sufficient to control ∼22000 genes in human cells. More importantly, we hypothesize, that this provides a way to coordinate multiple genes in a gene module to accomplish a particular biological function (*8–11*). However, due to the lack of experimental techniques, little is known about how TF combinations are organized to coordinate regulatory networks, which remains the central question in eukaryotic biology.

Most TF binding sites in eukaryotes have been identified by chromatin immunoprecipitation ChIP-on-chip (*12*) and ChIP-seq (*13, 14*), analyzing one TF at a time, through antibody pulling down followed by DNA sequencing, with a low spatial resolution of a few hundred base pairs. A common method for determining cellular binding motifs is footprinting, i.e. detecting DNA sequences that are protected from enzymatic treatment, most notably by DNase-seq, which detects the endpoints of DNA fragments cleaved by DNase I (*15*). However, both ChIP-seq and DNase-seq require bulk samples with millions of cells, usually of cell lines, and are incapable of detection on a single DNA.

To overcome these limitations, another footprinting method by cytosine methylation had been previously attempted with Single Molecule Footprinting (SMF) (*16*). However, it is not appliable to normal mammalian cells, as the extraneous methyltransferase used was mostly indistinguishable from endogenous methylation.

Here we report a novel approach based on double-stranded DNA cytosine deamination, named FOOtprinting with DeamInasE (FOODIE), to detect TFs’ binding sequences at single-base resolution along the genome, as the TFs’ steric hindrance prevents the conversion of cytosine to uracil. Cost-effective and scalable, FOODIE requires sequencing only thousands of cells, significantly fewer than those required for ChIP-seq or DNase-seq.

### Genome-wide mapping of TF footprints

FOODIE relies on double-stranded DNA deaminase with high efficiency (Fig. 1A). We initially tested DddA, the first double-stranded DNA deaminase reported (*17*), which only deaminates cytosine preceded by thymine, not capable of footprinting all TFs bound to the genome (Fig. S1A). To overcome this limitation, we characterized a new dsDNA cytosine deaminase from a large group of reported toxins (*18*), which we named DddB (also known as BadTF3 (*19*)) after DddA. Unlike DddA, DddB converts cytosine to uracil largely independent of adjacent bases (Fig. 1B), allowing us to footprint almost all TF motif sequences (Fig. S1A). We determined the crystal structure of DddB in complex with its immunity protein at 2.5 Å resolution (Fig. 1C). Despite the overall structural similarity between DddB and DddA, DddB features an intervening helix near the active site, whereas DddA has a loop in this region (Fig. S1B).

**Fig. 1.**
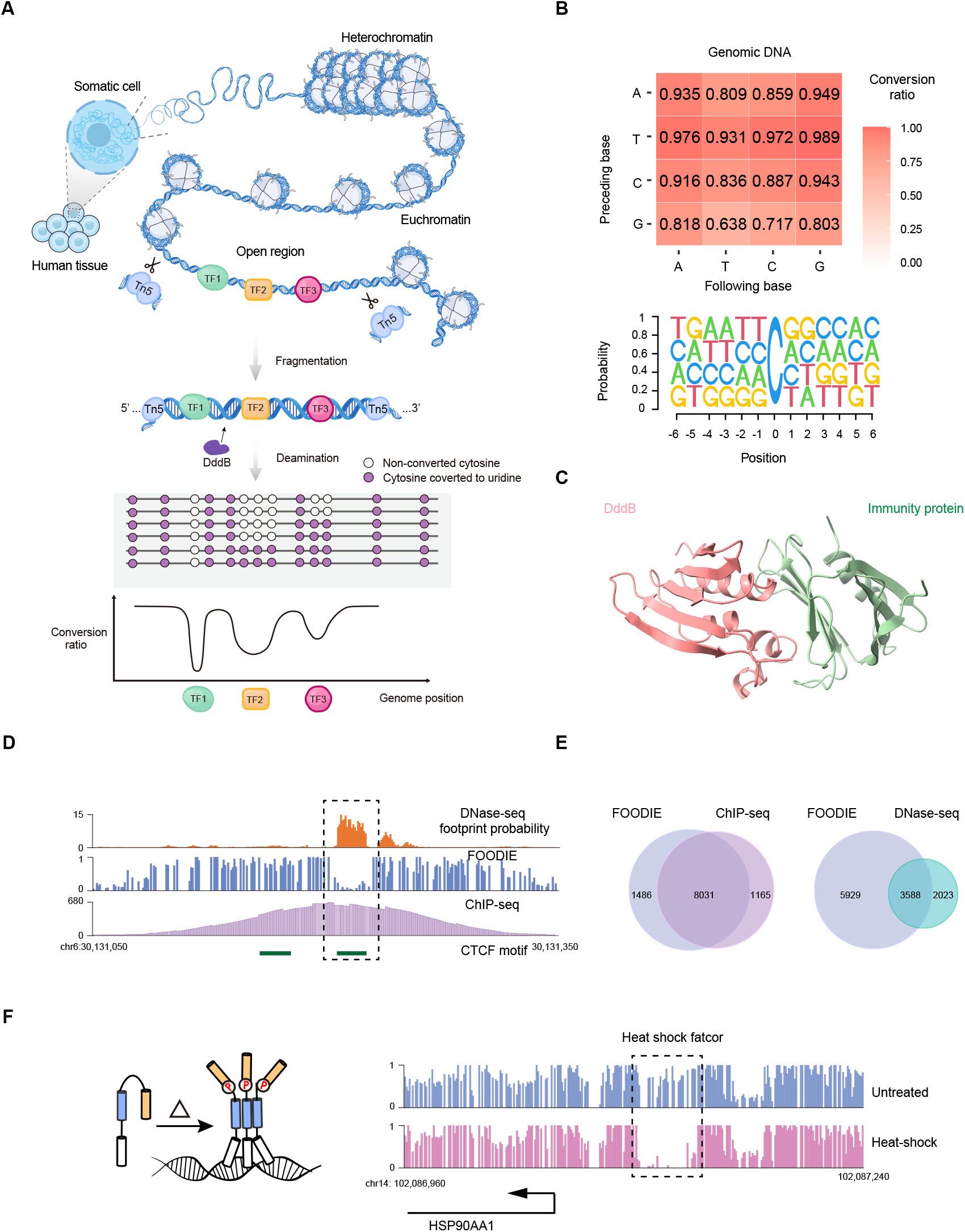
Genome-wide mapping of TF binding sites by FOODIE. (A) Workflow of FOODIE. Cells were permeabilized for Tn5 tagmentation and DddB deamination. Amplicons of DNA undergone *in situ* enzymatic deamination were sequenced to identify TF footprints. (B) (Top) DddB’s C to U conversion efficiency of naked DNA *in vitro.* The x-axis is the base following, and the y-axis base proceeding the cytidine. (Bottom) Sequence logo of the above data. (C) The crystal structure of DddB (red) complexed with its immunity protein (green). (D) Human genome tracks of DNase-seq footprint probability, FOODIE, and ChIP-seq in GM12878 cell line. The horizontal green bars represent the two possible CTCF motif sequences. Only the right one is acctually bound by CTCF. (E) Venn diagrams of FOODIE with ChIP-seq or DNase-seq for the numbers of CTCF binding sites detected in open chromatin regions of GM12878. (F) Heat shock factor (HSF) responding to heat shock. Upon heat shock, HSF monomers trimerize, bind to the Hsp90AA1 promoter, and leave a TF footprint on DNA.

Similar to but one step more than ATAC-seq (*20*), FOODIE uses transposase Tn5 in permeabilized cells to enrich the open chromatin regions, where most TFs bind and exert their regulatory roles. DddB is then added after Tn5 tagmentation to detect protein binding on DNA, whose underlying cytosines are protected from enzymatic deamination. As such, FOODIE is capable of probing the regulatory sites in open chromatin regions.

TF binding profiles detected by ChIP-seq, FOODIE, and DNase-seq are shown in Fig. 1D. ChIP-seq had a low spatial resolution (200-500bp), not capable of telling whether both or just one of the detected CTCF sequences were occupied. In contrast, FOODIE accurately identified CTCF that the footprint on the right is bound, which was consistent with DNase-seq footprinting. We further compared the number of genome-wide binding sites of CTCF detected by FOODIE with those obtained from ChIP-seq and DNase-seq. The CTCF binding sites detected by FOODIE show strong concordance with those detected by ChIP-seq and largely overlap with those detected by DNase-seq footprinting (Fig. 1E).

Next we employed a heat shock experiment to validate the reliability of FOODIE. Upon a temperature increase, the transcription factor HSF is activated to bind to the promoters of target genes to induce expression (*21*). At HSP90AA1 promoter, we observed a decrease in the conversion ratio within HSF motif sequences, which corresponded to the gain of HSF binding upon heat shock (Fig. 1F). The above data suggested that FOODIE can identify precise binding of TFs with high accuracy.

To support the interactive analyses of genome-wide footprints identified by FOODIE, we developed a web server with a user-friendly interface. This platform includes a genome browser for visualization and a data portal with our FOODIE data. Fig. 2A shows a browser display of the genomic region around the TRNAU1AP promoter in chromosome 1, a chromatin open region between nucleosomes -1 (and -2) and +1 (and +2, +3, +4), with three distinct footprints. The first row is the ATAC track seen by Tn5 tagmentation. The second row is the C to U conversion efficiency showing the footprints. The raw reads of single-molecule sequences below, with the converted cytosines on the Watson strand (red) and the Crick strand (green), show the binding fractions along the genome.

**Fig. 2.**
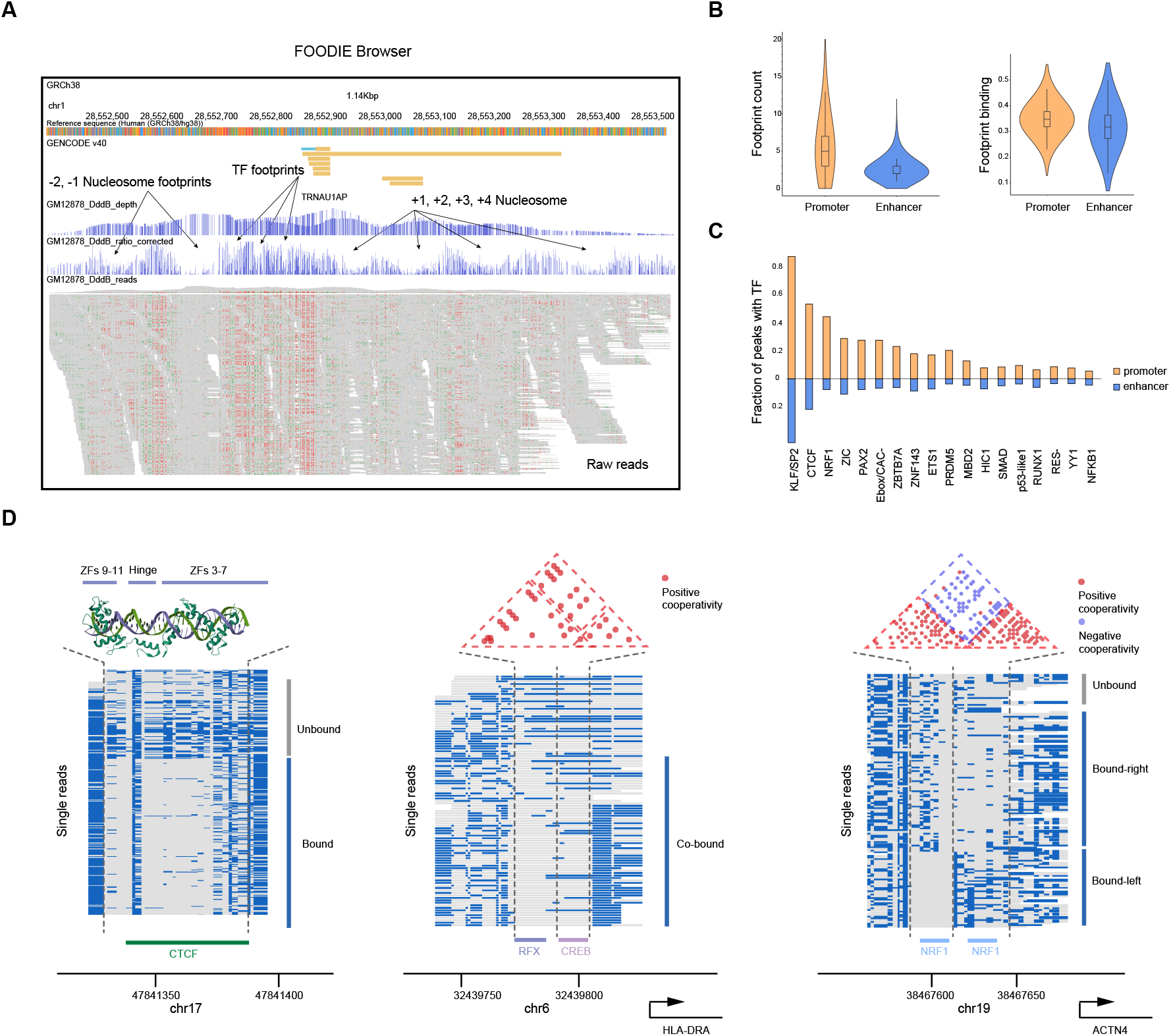
TF binding fractions and cooperativity among TFs detected by FOODIE. (A) FOODIE browser display at the promoter region of TRNAU1AP, with two nucleosomes at each side flanking the open chromatin region. The FOODIE sequencing depth is similar to ATAC-seq, whereas the FOODIE conversion ratio shows footprints of nucleosomes -1 & -2 and +1 & +2, and three distinct TFs in between. Below are single-molecule sequencing raw reads with the converted cytosines on the Watson strand (red) and the Crick strand (green). (B) (Left) Distributions for numbers of detected TF footprints in the promoters (median=5) and enhancers (median=2) in GM12878 cell line. (Right) Distributions for binding fractions of TF footprints in the promoters (median=0.348) and enhancers (median=0.317). (C) Abundance of the TF footprints with ChIP-seq-confirmed assignments on the promoters and enhancers. (D) (Left) Structure of CTCF (ZFs 3-11) and the single-molecule sequencing reads of FOODIE footprints on a CTCF binding site. (Middle) Single-molecule sequencing reads showing RFX and CREB footprints at the HLA-DRA promoter. The positive binding cooperativity between two TFs is evident from the positive Φ correlation coefficients (dots colored in red) of a cytosine pairs, each from either TF binding site. (Right) Single-molecule sequencing reads with two NRF1 binding sites at the ACTN4 promoter have negative binding cooperativity, which is evident from the negative Φ correlation coefficients (dots colored in blue) of cytosine pairs, each from either TF binding site.

We found that the deamination efficiency of DddB is dependent on the neighboring sequences of cytosines (especially from -4 to +2) (Fig. S2), which needs to be normalized when calculating the binding fraction. We measured deamination efficiency along the naked human genome as a positive control for the conversion ratio correction. The binding fraction is one minus conversion ratio.

Fig. 2B shows the mean number of footprints present in *cis*-regulatory elements of genes and their relative arrangement (see Material and Methods). The median number of footprints per gene promoter was found to be 5 (median number), compared to 2 in enhancer regions. Notably, the binding fractions of TFs in the promoter regions were higher than those of TFs in enhancer regions.

The FOODIE footprints need to be assigned to particular TFs, commonly through motifs tabulated on websites, such as JASPAR (*22*) and HOCOMOCO (*23*), which collected vast amounts of information from *in vitro* data of SELEX (*24*) and PBM’s (*25*) as well as *in vivo* data of ChIP-seq (*26*). It turned out that a footprint has many different possible assignments of TFs, which we can not pin down because of the lack of binding affinities from databases and the intracellular concentrations. While we are working on the methods to fully assign the footprints, in this work we assigned footprints to certain TFs only when they have strong ChIP-seq signal in GM12878 cell line. This resulted in reliable assignments to only seventy-nine known TFs.

As such, the abundance of the assigned TFs is shown in promoter and enhancer regions, we found promoters were particularly enriched for the NRF1, KLF/SP, ZNF143, ETS, and YY1, footprints, which were rarely detected in enhancers (Fig. 2C). Taken together, TF bound to promoters and enhancers differ in average number within open regions, binding fractions and TF types.

### Single-molecule sequencing reads revealing binding cooperativity among TFs

While the binding fraction of a particular binding site can be determined by the proportion of single molecules of sequenced DNA with unconverted C to that of the total of converted and unconverted C. The fact that sequencing occurs on individual molecules allows us to determine the cooperativity between TFs bound to two adjacent binding sites.

Single-molecule reads at a CTCF binding site in the genome are shown in Fig. 2D, exhibiting two separate footprints of the bridging structure of the zinc finger hinge (*27*), which are either simultaneously occupied or not. In Fig. 2D, two adjacent TFs RFX and CREB have two distinct footprints (*28*). To evaluate their binding cooperativity, we performed Fisher’s exact test for the significance of independence on a pair of cytosines, each selected from the two binding sites. The Φ correlation coefficient is between 0 and 1 for positive cooperativity and between -1 and 0 for negative cooperativity. A red dot indicates significant positive cooperativity (*P*<0.001). A pair of cytosines within the RFX (or CREB) binding sequence are positively cooperative because the binding of RFX to them is simultaneous. Interestingly, cytosine pairs across CREB and RFX binding sequences are also positively cooperative, i.e. RFX and CREB’s binding is not independent but facilitates each other (Fig. 2D). We attribute this positive coopertavity to protein-protein interactions between two TFs, which has been previously reported (*28*).

Conversely, negative cooperativity was seen between two NRF1 binding sites within the ACTN4 promoter (Fig. 2D). While the Φ of the two cytosines within each NRF1 binding site is positive, the pairs across different sites are negative (blue: *P*<0.001), meaning their binding is exclusive. Such single-molecule measurements provide unprecedented information about how TFs work together in the genome.

### Shared TFs among genes within the same CGM

The bursty nature of gene expression has been long established and attributed to the single-molecule nature of DNA in single cells (*29–31*). The coordination of a particular biological function by multiple genes requires their synchronization. People have been looking for modules of genes that working together for different biological functions, with the bulk (*11*) and single cells (*32*) by perturbation measurements. Notably, the synchronization of genes can also be probed by measuring the covariance matrix of any pair of mRNA or protein at single-cell levels. By improving the detectability of single-cell transcriptome and single-cell proteinome, we demonstrated the identification of correlated gene modules (CGMs) without perturbation under the steady state condition (*8, 9*).

Recently we have developed a new method for more accurate identification of CGMs, Triangle Screening of Hierarchical Accumulated-Edges (TriSHA) (*33*). Over Representation Analysis (ORA) reveals that nearly 80% of the modules are functionally annotated with a False Discovery Rate (FDR) below 0.05. Through module connection analyses, we have identified three distinct types of CGMs within the GM12878 cell line based on functional annotation, associated with specific cellular functions: housekeeping, tissue-specific, and cell cycle (Fig. 3A). Supplementary Table S1 lists the CGMs with size larger than 10 that we identified. Gene t-SNE plot of GM12878 clearly shows the cluster of genes in the identified CGMs (Fig. 3B).

**Fig. 3.**
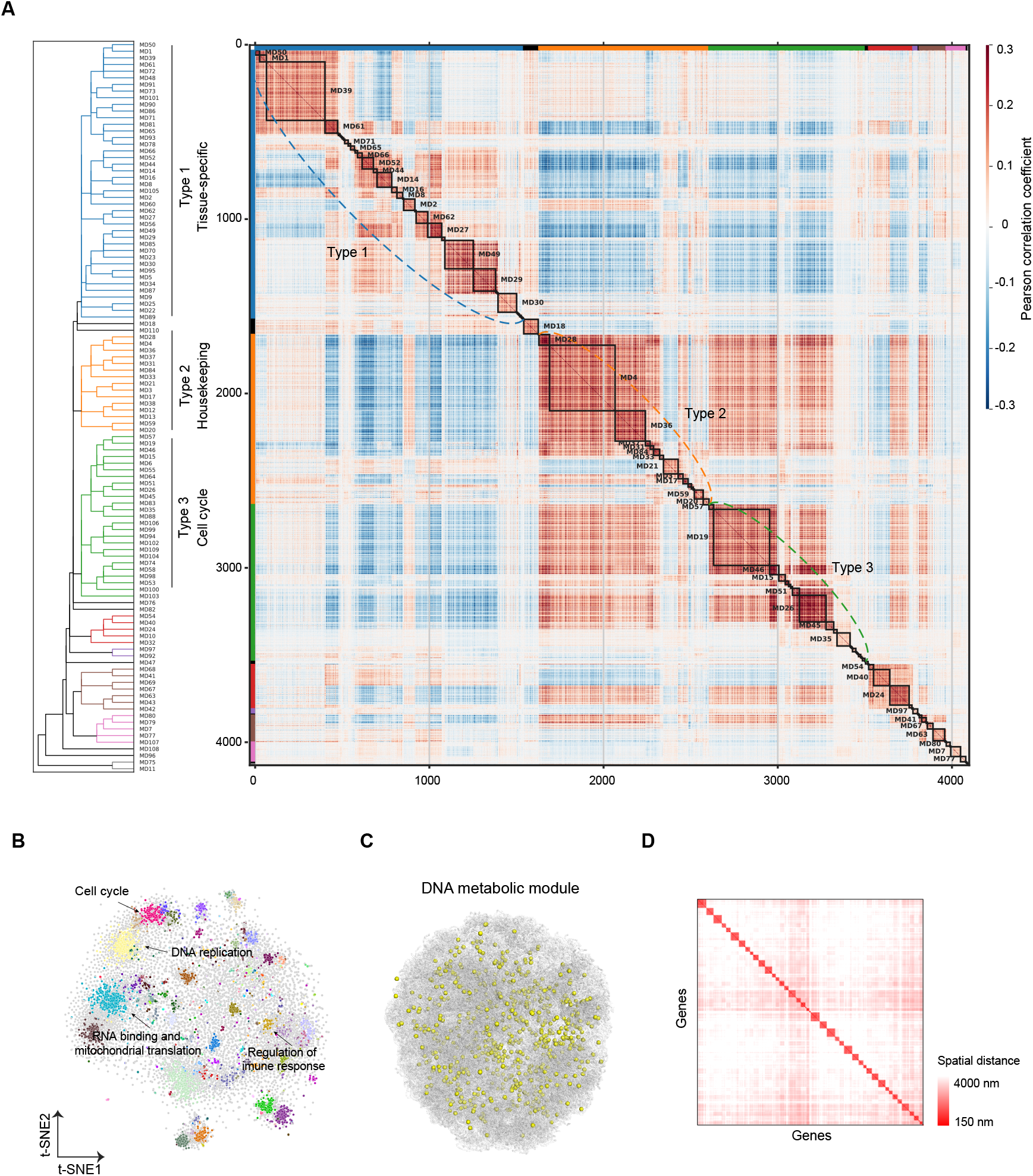
Genes in a CGM for a biological function not spatially clustered in the nucleus. (A) Pair-wise Pearson correlation coefficients for ∼ 4000 detected mRNA levels of GM12878 single cells under steady-state condition. CGMs were identified by the improved algorithm TriSHA (*33*), and indicated by enclosed squares along with the names of major CGMs. Shown on the left was the hierarchical clustering of CGMs with three major clades: cell cycle, housekeeping, and tissue-specific functions, respectively. (B) Gene t-SNE plot of the CGMs along with names of the representative ones. Each dot is an expressed gene, and genes in one CGM are labeled with a particular color. (C) 3D genome structure of a GM12878 cell experimentally determined by scMicro-C (*35*). Yellow points represent the promoter positions of the genes in the DNA replication CGM. (D) Pairwise spatial distance matrix for the genes in the DNA replication CGM, averaged over all scMicro-C structures, suggesting the genes in a CGM are evenly distributed in the nucleus rather than clustered.

How are the genes in the same module distributed within the 3D genome? Are they clustered together, or spatially dispersed? Our group determined the 3D genome structure of a somatic human cell (*34*). To answer the above question, we employed our latest high-resolution single-cell 3D genome structures of GM12878 cells using scMicro-C (*35*). Taking the DNA replication CGM (MD20, cell cycle classification) of 317 genes as an example, Fig. 3C shows that they are not clustered but relatively randomly distributed within the nucleus, with the more quantitative distribution of pair-wise distance shown in Fig. 3D. No spatial clustering of genes in other gene modules was observed either. This observation does not support the idea that the correlated genes are transcribed within certain locations in the nucleus.

We hypothesize that the coordination of different genes in the module is achieved through the binding of the same TFs in the contiguous euchromatin space. It has been reported that intranuclear TF levels in an eukaryotic cell often oscillates in time (*36, 37*). Multiple genes controlled by these TFs are thus expected to be synchronized.

With FOODIE, we are in a position to determine how TFs organize the CGMs. We looked at the enrichment of TFs shared among both promoters and enhancers of genes within a module, which is characterized by the hypergeometric test. Out of sixty-five CGMs identified in GM12878 (Fig. 3A and Supplementary Table S1), we successfully identified that many TFs are statistically enriched in different gene modules, including for example, E2F factors in the promoters of genes in MD16 cell cycle-related CGM (*38*) and RELB in the enhancers of genes in MD47 regulation of immune system CGM (*39*) (Fig. 4A). As a control, when we randomly selected tens of genes (size from 30 to 150) out of the ∼4000 expressed genes, no TFs were significantly enriched.

**Fig. 4.**
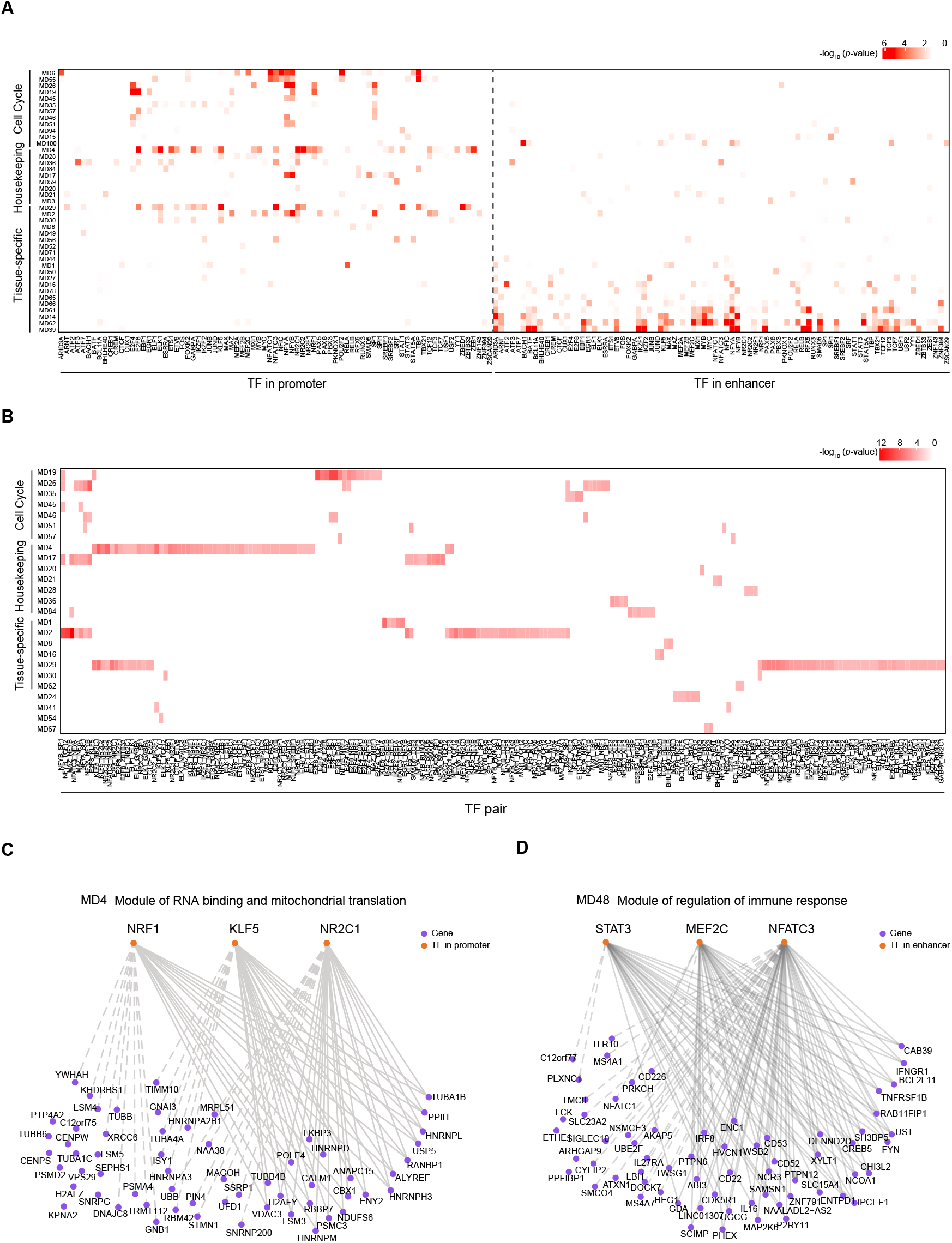
Genes in a CGM sharing TF(s) in their promoters and enhancers. (A) Heatmap showing the TF enrichment (-log *p*-value) within the major CGMs. Each row indicates a CGM and each column indicates TFs assigned by ChIP-seq to the measured footprints. The TF enrichments at promoters are shown in the left, and the TF enrichments at enhancers are on the right. Color density denotes the -log *p*-values calculated by the hypergeometric test. (B) Heatmap showing the TF pairs enrichment (-log *p*-value) within the major CGMs. Each row indicates a CGM and each column indicates TF pairs. Similarly, the -log *p*-values were calculated by the hypergeometric test. (C) NRF1-KLF5-NR2C1 TF combination at promoters regulating the genes in the M4 RNA binding and mitochondrial translation CGM. The solids lines are between a gene that connects more than one TFs, whereas the dash lines are between a gene with only one TF specified (and other un-assigned TFs). (D) STAT3-MEF2C-NFATC3 TF combination at enhancers regulating the genes in the MD48 regulation of immune response CGM.

Notably, we found enrichments of specific TFs are clearly different for the three types of CGMs (Fig. 4A). For cell cycle and housekeeping CGMs, the TF enrichment is primarily in promoters. For example, the NF-Y factor is enriched in the genes of MD14 cholesterol metabolism CGM. In contrast, for tissue-specific CGMs, the TF enrichment is primarily in enhancers. For example, STAT3 enrichment is present in MD 48 immune response CGM. This suggests that there are two distinct modes of CGM regulation: genes with essential cellular functions rely on TFs’ binding at promoters for robustness, while tissue-specific genes depend more on enhancers for larger tunability.

We further observed certain TF pairs are shared among genes in both promoters and enhancers within the same CGM (Fig. 4B) and evaluated the enrichment of TF using the hypergeometric test. TF pairs are highly abundant in GCMs, and there are more TF pairs in enhancer regions (Fig. S2A). Around one-third of the CGMs have enriched TF pairs, either at the promoters (Fig. 4B), or at the enhancers (Fig. S2B). A higher fraction is expected when TF footprints are fully assigned. This suggests the coordination of different genes in a module for a particular biology function through TF combinations is ubiquitous and by no means random. Concrete examples of TF pairs shared by multiple genes in a CGM are shown in Fig. 4C and 4D for promoters and enhancers.

As such, we are able to use FOODIE to probe how regulatory networks are orchestrated by a combination of TFs. Although we do not yet have all the TF footprints assigned, even at incomplete set (seventy-nine), already uncovered TFs pairs previously unknown in order to coordinate gene modules.

### Footprinting after cell typing by scFOODIE in heterogeneous tissues

Our goal is to systematicly characterize different human tissues with FOODIE and determine their genome-wide TF footprints. However, human tissues are heterogeneous with many different cell types, so it is necessary to determine each cell’s identity. We note that each gene only has one maternal allele and one paternal allele in an individual cell, which are not sufficient to provide reliable statistics for TF footprints. However, thousands of single cells identified for a specific cell type would be sufficient to carry out footprinting.

Similar to cell typing by single-cell transcriptome or single-cell ATAC-seq, we use FOODIE signal to do unsupervised clustering of a population of cells. We evaluated the “non-convertable ratio”, i. e. the ratio of unconverted cytosines to the total number of detected cytosines within the gene body ± 1000 bp regions for each gene. Using four well-characterized human cell lines (GM12878, K562, HEK293T, and HeLa) to generate a mix of single-cell FOODIE data, different cell types were clearly identified (Fig. 5A). After merging single-cell data of a particular cell type (GM12878 for example) as a bulk of pure population, we observed precise TF footprints of CTCF and NRF1 at the specific genome locations (Fig. 5A).

**Fig. 5.**
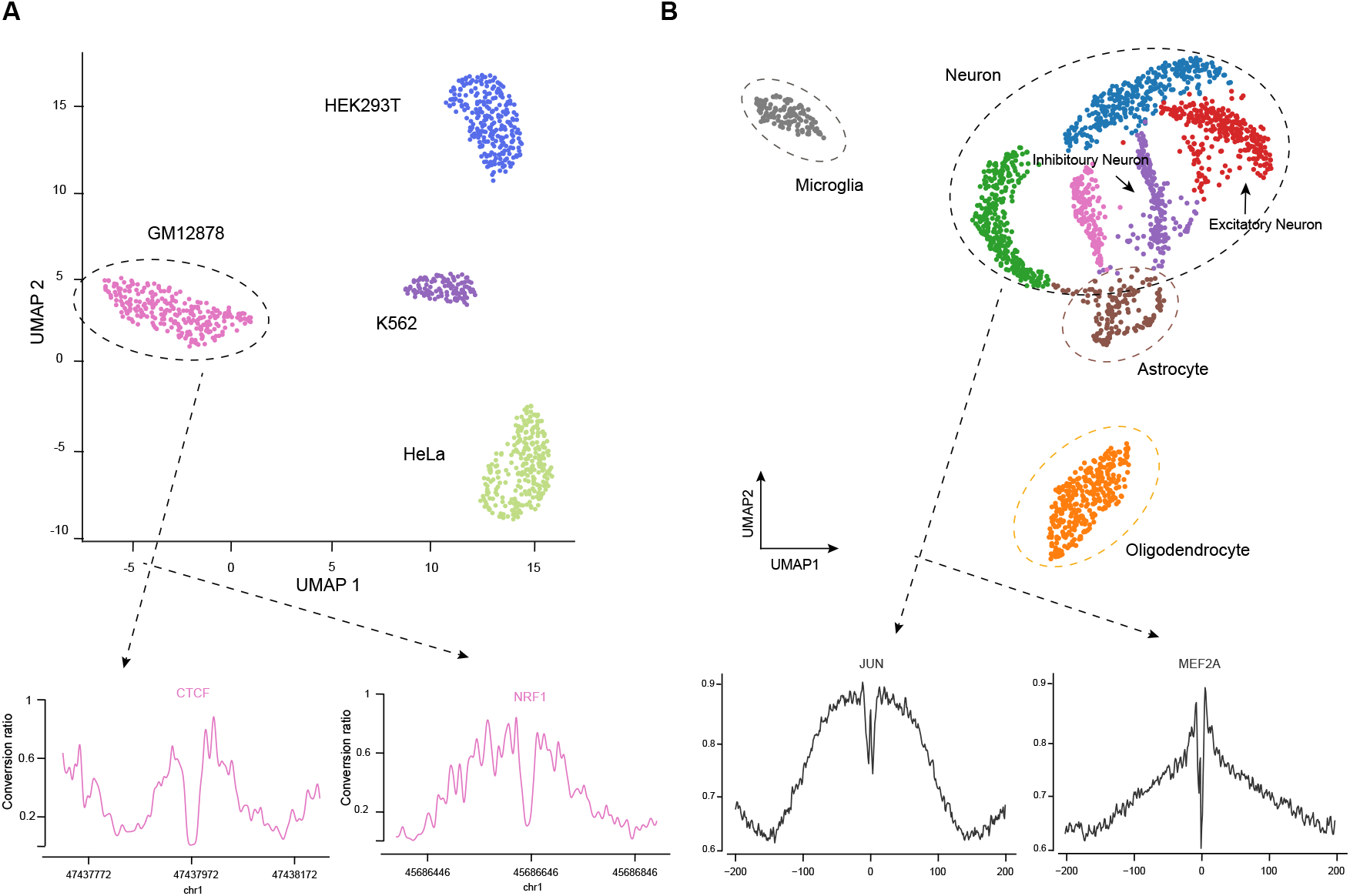
Cell-type-specific TF footprinting in heterogeneous tissues by scFOODIE. (A) Cells from a mixture of four human cell lines are classified into four different cell types based on their barcoded scFOODIE data. Unsupervised clustering was performed using UMAP. CTCF and NRF1 footprints at two genome locations from pooled cells of the GM12878 cell type are shown in the inset. (B) Cell typing of ∼1500 mouse hippocampus cells based on their scFOODIE data. Different colors indicate distinct cell types, astrocytes, microglia, oligodendrocytes, and neurons (including inhibitory and excitatory neurons), with annotations based on the marker genes. Footprints of JUN and MEF2B in neurons at merged genome-wide binding sites are shown in the inset.

We further conducted scFOODIE for ∼1500 cells from the mouse hippocampus. Four main cell types were identified, including oligodendrocytes, microglia, astrocytes, and neurons (both inhibitory and excitatory), which were annotated by marker genes (Fig. 5B). Using merged scFOODIE data of neurons, we observed footprints of JUN and MEF2B, two TFs critical for neuronal functions (Fig. 5B). Experiments like this are expected to provide new information of neuronal gene regulation in response to either physical or chemical stimulates in the brain.

### Outlook

In conclusion, we have shown that FOODIE is a high-precision method for genome-wide *in situ* mapping of TF footprints inside the nucleus with single-base resolution. FOODIE is experimentally simple, high-throughput, and cost-effective. ScFOODIE allows for cell typing in a heterogeneous tissue sample and TF footprinting in any cell type within a complex tissue. A systematic FOODIE database of different human tissues is underway. We anticipate that such information about TF combinations of different cell types and correlated gene modules will lead to the fundamental understanding of transcriptional regulation and cellular functions in eukaryotic biology, especially for human biology, which might hold the potential for understanding many diseases.

## Acknowledgments

We thank Prof. Ge Gao for the critical reading of the manuscript. We thank Qiuwan Liu for helping with the figures, and Wenzhong Zhang for helping with the web server.

## Author contributions

X. S. X., R. H., W. D., Z. W., and W. M. designed the experiments. R. H., W. D., W. M., Z. W., Y. Y., Y. Z., S. L., C. L., and J. R. collected the experimental data. L. G., C. X., D. L., K. S., F. J., J. Z., X. W., Y. P., Z. Zhang, X. C. and Z. Zhu performed the data analyses. X. S. X. supervised the project and wrote the manuscript with the help of all authors.

## Materials and Methods

Materials and methods are placed in the supplementery document.

## Supplementary Materials

**Figure S1.**
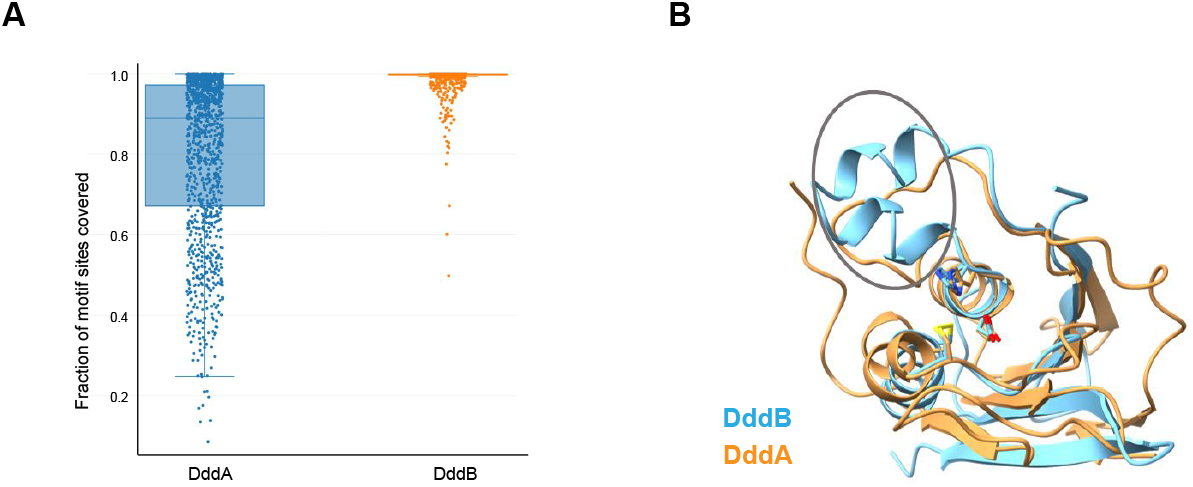
(A) Percentage of TF motifs on the human genome that can be assessed by DddA (probing every TC) and DddB (probing every C) enzymes. (B) Comparsion of cystal sturcutes of DddA and DddB.

**Figure S2.**
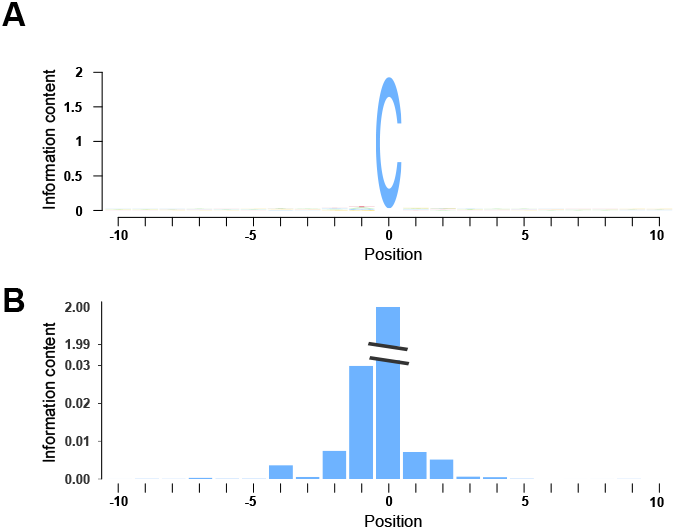
(A) Sequence logo of DddB conversion yields for a cytosine on naked DNA *in vitro*. (B) Bases from -4 to +2 positions of cytosine show relatively large bias on conversion.

**Figure S3.**
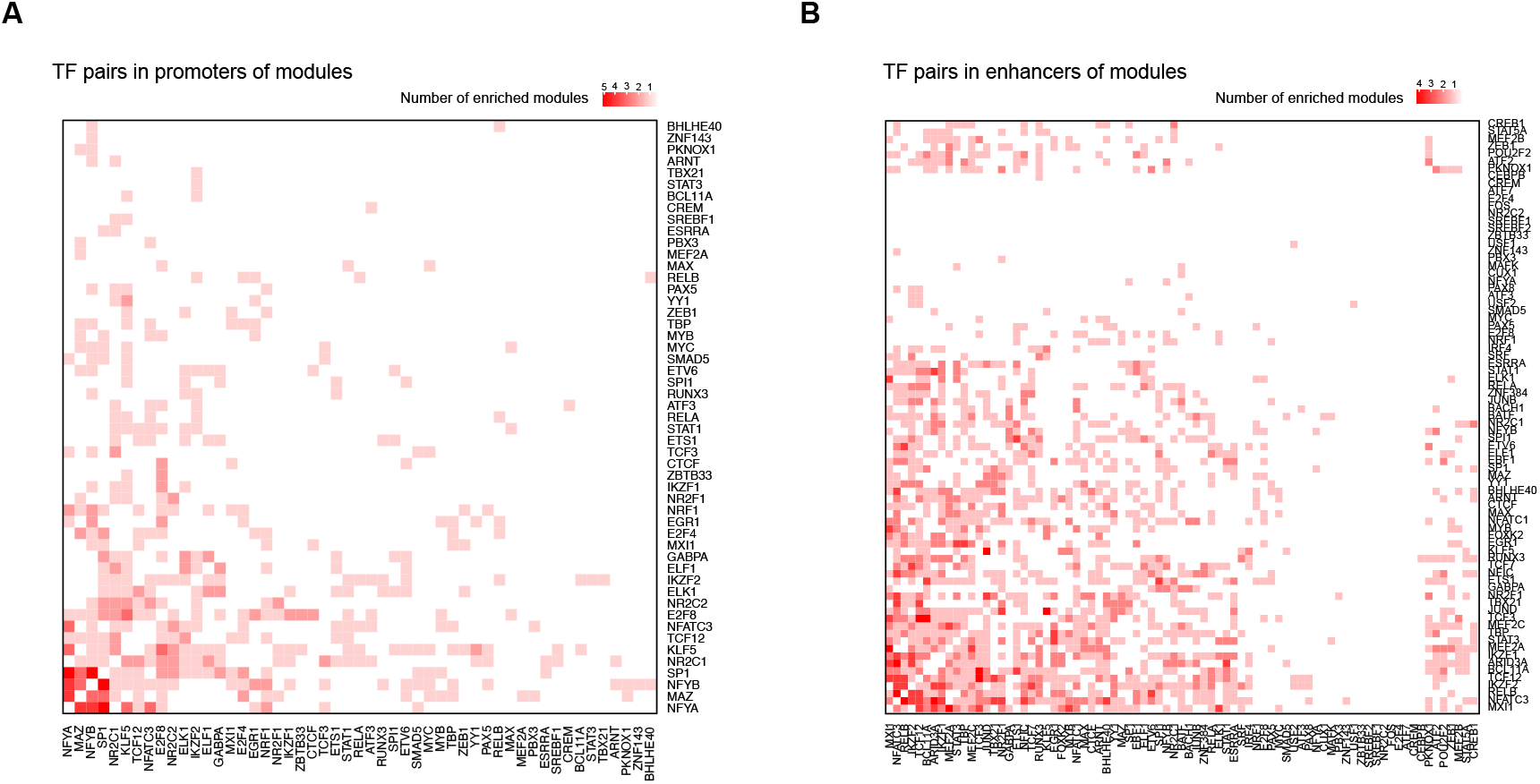
Enhancers harbor more enriched TF pairs than promoters. (A) Heatmap showing the number of modules, in which the TF pairs in a promoter are enriched. Color bar on the top denotes the number of enriched modules. (B) Heatmap showing the number of modules, in which the TF pairs in a enhancer are enriched. Color bar on the top denotes the number of enriched modules.

