## Supplementary material for "Human transcription factor combinations mapped by footprinting with deaminase": Table S1

| module | group | module_code | module_size |
|---|---|---|---|
| MD1 | tissue_specific | 2_0 | 39 |
| MD2 | tissue_specific | 2_1 | 65 |
| MD3 | house_keeping | 2_2 | 28 |
| MD4 | house_keeping | 2_3 | 376 |
| MD5 | cell_cycle | 2_5 | 19 |
| MD6 | Unknown | 2_6 | 56 |
| MD7 | tissue_specific | 2_7 | 33 |
| MD10 | tissue_specific | 2_13 | 84 |
| MD11 | cell_cycle | 2_14 | 37 |
| MD13 | tissue_specific | 3_0 | 30 |
| MD14 | house_keeping | 3_1 | 29 |
| MD15 | Unknown | 3_2 | 83 |
| MD16 | cell_cycle | 3_3 | 321 |
| MD17 | house_keeping | 3_4 | 33 |
| MD18 | house_keeping | 3_5 | 84 |
| MD19 | house_keeping | 3_8 | 111 |
| MD20 | cell_cycle | 4_0 | 153 |
| MD21 | tissue_specific | 4_1 | 80 |
| MD22 | house_keeping | 4_2 | 61 |
| MD23 | tissue_specific | 4_3 | 128 |
| MD24 | tissue_specific | 4_4 | 107 |
| MD27 | cell_cycle | 4_9 | 75 |
| MD28 | house_keeping | 5_0 | 174 |
| MD31 | tissue_specific | 6_0 | 336 |
| MD32 | house_keeping | 6_1 | 93 |
| MD33 | Unknown | 6_2 | 26 |
| MD34 | tissue_specific | 6_5 | 21 |
| MD35 | cell_cycle | 6_6 | 44 |
| MD36 | cell_cycle | 6_7 | 53 |
| MD37 | tissue_specific | 6_10 | 163 |
| MD38 | tissue_specific | 7_0 | 26 |
| MD39 | cell_cycle | 7_1 | 40 |
| MD40 | tissue_specific | 7_2 | 65 |
| MD41 | house_keeping | 7_4 | 28 |
| MD42 | cell_cycle | 7_5 | 12 |
| MD43 | tissue_specific | 7_6 | 18 |
| MD44 | cell_cycle | 9_0 | 27 |
| MD46 | house_keeping | 9_2 | 51 |
| MD47 | tissue_specific | 10_0 | 73 |
| MD48 | tissue_specific | 10_1 | 67 |
| MD49 | Unknown | 10_2 | 67 |
| MD51 | tissue_specific | 13_0 | 22 |
| MD52 | tissue_specific | 13_1 | 26 |
| MD53 | Unknown | 13_2 | 32 |
| MD54 | Unknown | 14_0 | 14 |
| MD55 | tissue_specific | 17_0 | 20 |
| MD57 | Unknown | 22_0 | 30 |
| MD58 | tissue_specific | 24_0 | 14 |
| MD62 | house_keeping | 30_0 | 35 |
| MD63 | cell_cycle | 55_0 | 19 |
| MD64 | Unknown | 67_0 | 27 |
| MD65 | cell_cycle | 84_0 | 14 |

| **GO_annotation** |
| --- |
| MD1 GOBP_RESPONSE_TO_CYTOKINE,WP_TNFRELATED_WEAK_INDUCER_OF_APOPTOSIS_TW |
| MD2 GOCC_ENDOPLASMIC_RETICULUM,GOCC_ENDOPLASMIC_RETICULUM_CHAPERONE_CO |
| MD3 WP_GLYCOLYSIS_AND_GLUCONEOGENESIS,WP_AEROBIC_GLYCOLYSIS,GOBP_ORGANIC_ |
| MD4 GOMF_RNA_BINDING,REACTOME_MITOCHONDRIAL_TRANSLATION |
| MD5 GOCC_DNA_PACKAGING_COMPLEX |
| MD7 GOMF_TAP_BINDING,GOBP_ANTIGEN_PROCESSING_AND_PRESENTATION_OF_ENDOGEN |
| MD10 GOCC_ORGANELLE_SUBCOMPARTMENT,GOBP_ESTABLISHMENT_OF_PROTEIN_LOCALI |
| MD11 GOBP_RNA_PROCESSING |
| MD13 GOCC_MHC_CLASS_II_PROTEIN_COMPLEX |
| MD14 WP_CHOLESTEROL_METABOLISM_WITH_BLOCH_AND_KANDUTSCHRUSSELL_PATHWA |
| MD15 WP_P53_TRANSCRIPTIONAL_GENE_NETWORK,GOBP_CELLULAR_RESPONSE_TO_STRESS |
| MD16 GOBP_DNA_METABOLIC_PROCESS,GOCC_CHROMOSOME,REACTOME_DNA_STRAND_EL |
| MD17 KEGG_HUNTINGTONS_DISEASE,GOCC_MEMBRANE_PROTEIN_COMPLEX,GOCC_RESPIRA |
| MD18 GOBP_RESPONSE_TO_OXYGEN_LEVELS,WP_PHOTODYNAMIC_THERAPYINDUCED_HIF1_ |
| MD19 GOMF_RNA_BINDING,GOBP_RNA_LOCALIZATION |
| MD20 GOBP_CELL_CYCLE,GOBP_MITOTIC_SISTER_CHROMATID_SEGREGATION |
| MD21 GOBP_DEFENSE_RESPONSE_TO_SYMBIONT,GOBP_BIOLOGICAL_PROCESS_INVOLVED_I |
| MD22 REACTOME_MRNA_SPLICING,GOMF_RNA_BINDING,REACTOME_THE_ROLE_OF_GTSE1_ |
| MD23 REACTOME_EUKARYOTIC_TRANSLATION_ELONGATION,KEGG_RIBOSOME,REACTOME_ |
| MD24 GOCC_CATALYTIC_COMPLEX,REACTOME_EUKARYOTIC_TRANSLATION_ELONGATION |
| MD28 GOBP_RIBOSOME_BIOGENESIS,GOMF_RNA_BINDING,GOCC_PRERIBOSOME |
| MD31 GOBP_ACTIN_FILAMENT_BASED_PROCESS,GOBP_LOCOMOTION,GOCC_CELL_SUBSTRA |
| MD32 GOCC_GOLGI_APPARATUS,GOBP_REGULATION_OF_ADAPTIVE_IMMUNE_RESPONSE |
| MD33 GOMF_CATALYTIC_ACTIVITY_ACTING_ON_A_NUCLEIC_ACID,GOCC_RIBONUCLEOPROT |
| MD35 GOCC_CENTROSOME,GOCC_MICROTUBULE_CYTOSKELETON,PID_FOXM1_PATHWAY |
| MD36 REACTOME_CELL_CYCLE,GOBP_CELL_CYCLE |
| MD38 GOBP_IMMUNE_SYSTEM_DEVELOPMENT,GOBP_T_CELL_DIFFERENTIATION |
| MD39 GOBP_MITOTIC_CELL_CYCLE_PROCESS,GOBP_CELL_CYCLE,PID_PLK1_PATHWAY |
| MD40 GOBP_REGULATION_OF_INTRACELLULAR_SIGNAL_TRANSDUCTION,GOBP_VACUOLE_O |
| MD41 GOCC_MITOCHONDRION,GOCC_INTRACELLULAR_PROTEIN_CONTAINING_COMPLEX |
| MD42 REACTOME_DNA_DAMAGE_TELOMERE_STRESS_INDUCED_SENESCENCE,REACTOME_C |
| MD43 GOMF_CYTOKINE_ACTIVITY,GOBP_CELLULAR_RESPONSE_TO_INTERLEUKIN_1,GOBP_L |
| MD44 REACTOME_MITOTIC_G1_PHASE_AND_G1_S_TRANSITION,PID_E2F_PATHWAY,GOBP_CE |
| MD46 GOBP_ACTIN_FILAMENT_BASED_PROCESS,GOBP_CYTOSKELETON_ORGANIZATION,REA |
| MD47 GOBP_REGULATION_OF_IMMUNE_SYSTEM_PROCESS,GOBP_LEUKOCYTE_CELL_CELL_A |
| MD48 GOBP_REGULATION_OF_IMMUNE_RESPONSE,GOBP_IMMUNE_RESPONSE,GOBP_REGULA |
| MD49 GOBP_NITROGEN_COMPOUND_TRANSPORT,GOBP_GOLGI_VESICLE_TRANSPORT |
| MD51 REACTOME_SIGNALING_BY_RHO_GTPASES_MIRO_GTPASES_AND_RHOBTB3,REACTOM |
| MD52 GOBP_REGULATION_OF_CELL_DEATH,GOBP_TRANSMEMBRANE_RECEPTOR_PROTEIN_ |
| MD53 GOBP_CELLULAR_AMINO_ACID_METABOLIC_PROCESS,GOBP_SMALL_MOLECULE_MET |
| MD55 GOMF_LIPID_BINDING,GOBP_CELL_MIGRATION |
| MD62 GOCC_INNER_MITOCHONDRIAL_MEMBRANE_PROTEIN_COMPLEX,GOCC_MITOCHONDR |
| MD63 KEGG_WNT_SIGNALING_PATHWAY,GOCC_CHROMOSOME |
| MD65 GOBP_PEPTIDYL_AMINO_ACID_MODIFICATION |

| module_genes |
| --- |
| TNFAIP3,ICAM1,NFKBIA,BIRC3,NFKBIE,SNX11,FAS,RELB,CD80,NECAP2,DEPP1,SERPINB8, |
| HSPA5,HSP90B1,MANF,DNAJB11,CALR,SDF2L1,PDIA6,CANX,PDIA3,PPIB,SPCS3,AC092069.1, |
| TPI1,ENO1,PGAM1,LDHA,PGK1,GPI,PKM,GAPDH,PFKP,FAM162A,DARS,HK2,ENO2,HK1,PGA |
| CCT5,HSPD1,TCP1,HSP90AA1,CCT4,HSP90AB1,HSPE1,CCT2,CCT6A,RAN,CYCS,HSPH1,HSPA |
| HIST1H2AC,HIST1H2BJ,HIST1H4H,HIST1H1C,HIST1H2BG,HIST1H2BK,HIST2H2BE,HIST1H2B |
| AC006547.3,PEAK1,RUBCN,AC011558.1,AC073641.1,ORMDL2,ETFDH,MKLN1-AS,ENOSF1,CA |
| HLA-C,HLA-A,HLA-B,B2M,PSMB9,OGA,TAPBP,LAPTM5,TAP1,CCNDBP1,HLA-F,TENT5A,FA |
| IGLC3,IGHM,IGHV3-23,IGLC2,SUB1,UBE2J1,IGLV6-57,SSR3,TNFRSF17,IGLV2-5,SEC62,MTDH |
| AC109326.1,AC007952.4,SNORD3B-1,SNORD3B-2,AC245014.3,RNVU1-19,AL021155.2,RNVU1-1 |
| HLA-DQA1,HLA-DRA,HLA-DPB1,HLA-DQB1,CD74,HLA-DRB1,HLA-DPA1,HLA-DOB,HLA-DM |
| MSMO1,IDI1,HMGCS1,HMGCR,SQLE,INSIG1,FDFT1,SCD,DHCR24,ACAT2,LDLR,DHCR7,SC5D |
| TM7SF3,RPS27L,CMBL,DDB2,CDKN1A,MDM2,GADD45A,SPATA18,XPC,ZMAT3,FUCA1,TRI |
| MCM3,MCM4,MCM6,PCNA,NASP,GINS2,MCM2,FEN1,MCM7,HELLS,CLSPN,CHEK1,DNMT1, |
| CALM3,UBL5,NEDD8,MYL6,COX6C,TMSB10,COX7A2,HADHA,PSME1,SRP14,NDUFA4,VAMP |
| APOL1,KDM3A,P4HA1,ZNF395,PFKFB4,BNIP3L,ALDOC,ZNF292,FUT11,PDK1,BNIP3,ZNF160, |
| ILF2,DHX9,CCT3,CACYBP,HNRNPU,SNRPE,RBM8A,TPR,EPRS,PRRC2C,UBAP2L,PARP1,PFD |
| CCNA2,TOP2A,NCAPG,SPAG5,NDC80,KIF11,FOXM1,CENPF,KIF4A,TPX2,MKI67,AURKB,HM |
| TLR7,FCRL5,MX1,OAS1,EPSTI1,OAS2,XAF1,IFI44,IFI44L,SP110,STAT1,IFIT1,SAMD9,SP100,SA |
| TUBBP1,KPNA2,TUBB,TUBA1B,TUBB4B,RANBP1,TUBA1C,SSRP1,HNRNPD,HNRNPA3,HNRN |
| RPS18,RPL3,RPL37A,RPS3A,RPS6,RPL11,RPS3,RPL5,RPL9,RPS4X,RPS20,RPL10A,RPL4,RPL7,R |
| EEF1D,RPL18A,RPL18AP3,RPS14,AL031727.1,UBXN1,AL365357.1,EEF1DP1,AL591846.1,NARF, |
| RNU4-39P,AC099778.1,AL031733.2,CSKMT,SREBF2-AS1,RN7SL832P,AP003733.4,AC067945.3,A |
| NOLC1,DDX21,NCL,NOP56,DKC1,NOP16,NOP58,MRTO4,EIF3B,GNL3,LYAR,SRSF7,RPF2,SYN |
| ANKRD33B,CCL22,CD86,CD44,UPB1,SRGN,CLIC2,DENND5A,TCF7,RASSF4,TGM5,SSTR2,KIF |
| MR1,MCL1,KCNN3,SLAMF7,BTG2,ASH1L,RCSD1,ARID4B,WDR26,CTSS,CD46,TRAF3IP3,AL3 |
| USP10,VPS35,BRD7,CNOT1,FAM192A,NFATC3,NOB1,DNAJA2,TERF2,COQ9,CBFB,CMC2,BRD |
| ENAM,ITGA4,JCHAIN,RNF103,GNG7,TXNIP,DENND5B,RUFY3,COL24A1,OXR1,AC012368.1,L |
| CCNB1,CDC20P1,CDC20,CCNB2,PRR11,DLGAP5,NEK2,CDKN3,HMMR,PLK1,PTTG1,CDC25B, |
| RRM2,RRM1,SMC2,ZWINT,DHFR,FANCI,CDCA5,MELK,ESCO2,POLQ,DIAPH3,RAD54L,STIL, |
| RPL9P7,RPS3AP6,RPS3AP26,RPL4P4,AC073861.1,RPS3AP47,RPS20P2,RPL9P9,EEF1A1,AC0080 |
| BCL2A1,CD40,CD83,TRAF1,PLEK,MARCKS,MKNK2,WDR91,NFKB1,TNIP2,MCOLN2,IL4I1,PL |
| SGO2,ASPM,KIF14,GTSE1,AURKA,CENPA,DEPDC1,KIF20A,KNSTRN,BORA,GPSM2,ECT2,UB |
| KLHL24,TP53INP1,YPEL5,TMEM140,PNRC1,HBP1,CREBRF,PCMTD1,FAM117A,PIM2,SERINC |
| NDUFS2,UFC1,COPA,ENSA,PSMB4,SDHC,RNPEP,POLR3GL,C1orf43,DCAF6,PSMD4,LAMC1,P |
| HIST1H4C,HIST1H2AE,HIST1H1D,HIST1H2BF,HIST1H2AJ,HIST1H1B,HIST2H2AC,HIST1H1E,H |
| CCL4,CCL3,DUSP2,CCL3L1,EGR2,IER2,TNF,EGR1,CD69,PHACTR1,ARL5B,XCL1,NAB2,SPAG |
| DTL,CDC25A,MCM10,CDC6,UNG,CCNE1,XRCC2,CCNE2,E2F2,ZGRF1,PSMC3IP,ZNF367,AUNI |
| TPM4,WDR1,LCP1,SRI,ACTB,AC098614.1,ARPC2,ACTR3,CAP1,MYL12A,ACTR2,TMSB4X,ACT |
| CFLAR,CACNA1E,IRF2BP2,MAL,SLAMF1,NCALD,NEK6,LSP1,STX6,THEMIS2,INSR,CCR7,SY |
| FYN,CD53,IFNGR1,CD22,PTPN6,PIK3AP1,SCIMP,IL27RA,MS4A1,TNFRSF1B,RAB11FIP1,UGC |
| AP1G1,N4BP1,COX4I1,TERF2IP,GLG1,CHD9,HERPUD1,IST1,MAP1LC3B,ANKRD11,GSE1,DYN |
| PFN2,FRY,TBC1D30,PTPN13,TSPAN12,B3GNT2,ARHGAP5,CTSC,AC104793.1,SEPT10,STARD1 |
| ST6GAL1,RAB30,RASSF6,RAPGEF2,SAMD12,SETX,ZDHHC14,FRMD4B,OGT,NLRP1,CDK13,R |
| PSAT1,SLC7A11,GPT2,ATF4,PCK2,CTH,PHGDH,CARS,SARS,WARS,ATF5,SLC3A2,PSPH,NFE2 |
| GOT2,CYB5B,NUDT21,KARS,NAE1,SF3B3,HSBP1,NUTF2,TMEM208,MLKL,GAN,TANGO6,E2 |
| CRYM,BMS1P8,CHL1,TBX15,TINAG,CCDC198,MYO6,ANXA1,CPNE3,PDE1C,CD101,CDCP1,L |
| SLC35B2,RF00019,MYH15,PLLP,RNU6-942P,CORO7,GTF2IP20,AC096992.2,FOXK1,AP000347.1, |
| MAP3K5,SYNE2,SIPA1L1,IKZF2,KIAA0319L,HERC2,LZTFL1,MBNL3,KCNQ5,USP48,MIB1,RB |
| HNRNPA1,PPA1,CHCHD2,NAP1L1,LDHB,SNRPD2,AHCY,PPIA,FBL,TXN,TMA7,SEM1,NDUFS |
| NIPBL,SPTBN1,ROCK1,SCAF11,CREBBP,PPP1R12A,CBL,ARHGEF12,FRYL,ZNF148,CHD1,APC |
| ANKRD36B,AC092683.1,LINC00342,ANKRD36,ANKRD36C,ZNF37BP,AC138409.2,AC138866.1, |
| MYB,RNF157,ARID2,CMAHP,UHRF2,SLAIN1,SGMS1,UNC13C,PRKRA-AS1,MANBAL,C6orf10 |

MTMR9,TRAF3,RAB21,POGLUT1,TANK,TNFAIP8,ZNF267,DENND3,DENND4A,HIVEP1,STARD4
DNAJC3,PDIA4,HYOU1,CRELD2,RPN1,SPCS2,PDIA3P1,MLEC,ATP2A2,ATP6V1C2,SSR1,MAGT
M1P8,AC006064.4,CXCR4,ADPRHL2,MIF-AS1,NCKIPSD,PRPSAP2,NDUFC1,CNOT11,GSTO1,GA
9,EBNA1BP2,HSP90AA2P,HSPA8,SNRPD1,PSMD1,NME1,SSBP1,STRAP,C1QBP,ATIC,HNRNPAB
C,HIST1H2BD,AL021807.1,AL031777.3,HIST2H2BF,HIST1H3G,HIST1H2BH,HIST2H4A,HIST1H3
CNA1A,RNF165,AC034229.1,CES2,INGX,CARMN,PIPOX,AC105429.1,LINC02141,TRRAP,TENM3
M49A,TMEM30A,CTSH,AC104699.1,SERTAD2,ZNF226,RBM5,IP6K2,ATP6AP1,ATP6V0E1,YIPF3
,MAN2A1,XBP1,IGHJ6,SCARB2,GALNT1,STRBP,B4GALT3,TRIB1,RHOQ,PRDM1,EIF2AK3,MEI
5,SNORD3A,RF00012,RN7SL2,SNORD3C,HIST4H4,VMP1,RNVU1-7,HIST1H4B,SNORD13,SNOR
A,HLA-DOA,HLA-DMB,CIITA,IL2RG,HLA-DPB2,S1PR2,HLA-DRB5,HCP5,IGFBP4,P4HA2,CYB5
),RDH11,FADS1,PANK3,IDH1,HSD17B7,NSDHL,ERG28,FADS2,MMAB,AL691447.2,LSS,TMEM9
I22,RRM2B,PURPL,TRIAP1,FDXR,AL158206.1,SESN2,CCP110,OTP,BLOC1S2,ASCC3,FBXO22,TI
RFC4,HNRNPF,MSH2,ATAD2,BRCA1,DNAJC9,RBBP8,SLBP,FAM111B,MCM8,ATAD5,WDHD1,V
8,NRDC,COX6B1,CTNNA1,LGALS1,CAPNS1,CHMP2A,POLR2G,AP2M1,CHP1,CD63,UROD,CNP
C4orf3,EGLN1,INSIG2,HILPDA,WSB1,AP3S1,VEGFA,SLC25A36,RNASET2,PLEKHA2,DDIT4,AC0
N2,TPM3,TOMM20,MRPL24,HDGF,NUCKS1,IFI16,IARS2,TIMM17A,HEATR1,ANP32E,IPO9,GNPA
GB2,KIF20B,SMC4,KIF15,ARHGAP11A,KIF2C,PBK,BUB1B,CDCA2,KNL1,ANLN,HJURP,CDK1,K
MD9L,IFI6,ITGB7,LRRK2,PARP15,DDX60,EIF2AK2,PARP9,U62317.4,OAS3,TRANK1,PTEN,PDL
IPA2B1,YWHAH,VDAC3,H2AFZ,SNRPG,PPIH,TUBA4A,PSMA4,PSMC3,XRCC6,HNRNPM,H2AF
RPL13A,RPS12,RPLP0,RPL14,RPSA,RPS2,RPL8,RPS16,RPS23,RPS11,RPL31,RPL27A,RPL6,RPS15A
RPL28,RPS15,AC007969.1,RHOC,PPP2R1A,HCLS1,IFI35,HEXA,HLA-E,FAU,TBCD,FLII,ARPC4,N
C098818.2,AC145285.3,AL512408.1,AL356488.3,NSMCE1-DT,AC068338.2,AC099343.3,AC011611
CRIP,NIFK,GPATCH4,WDR43,BRIX1,LARP1,TSR1,NOP56P1,NAA15,CEBPZ,ABCE1,PUM3,PAK1
26B,MARCKSL1,BASP1,FMNL3,SOX9,CD82,ARHGAP31,IL1R1,ENPP2,IL1R2,CRYZ,DTX4,RCN1
90728.4,MDM4,CEP350,KDM5B,SDCCAG8,UHMK1,ARHGEF2,CEP170,ELK4,LYPLAL1,CD48,RC
07P2,C16orf87,DDX19B,ADAT1,TENT4B,OGFOD1,TXNL4B,DDX19A,PDP2,CIAPIN1,AFG3L1P,D
INC01934,FNDC3B,METTL7A,AC243960.1,CFAP54,IFNG-AS1,DGKD,LINC00476,AIG1,ENDOV
ARL6IP1,TNFAIP8L1,TRMU,RNF5,LRRFIP1,PSMD10,CEP70,SAPCD2,JPT1,HP1BP3,DDX6,GIHCG
CDC45,NET1,NCAPG2,FANCD2,SKA3,FAM111A,PCLAF,RAD51AP1,UBE2T,MND1,NSD2,NCAPD
38.1,RPL7P9,RPL13AP5,AP001324.1,RPS23P8,Z74021.1,AC079140.2,RPL3P4,RPL6P27,AC026403.1
A1A,SWAP70,SPIB,RAB8B,RASSF2,STAT6,TET3,TESPA1,MAB21L3,RASGRP1,CNDP2,CXorf38,
E2C,FAM72D,ARHGAP19,FAM72C,FAM72B,CDC25C,G2E3,BRD8,PIF1,CDC27,CALM2,PNRC2,R
1,RIPOR2,SECISBP2L,FAM214A,GOLGB1,BAZ2B,GCC2,N4BP2L2,NCOA3,RICTOR,CLK1,NBR1,
DE4DIP,MGST3,DEDD,MRPS14,LAMTOR2,TMCO1,TMEM9,DEGS1,INAVA,LIX1L,LINC00467,TS

P,USP37,ZNF100,ZNF519,E2F3,ZMYND19,GJC1,CCDC138,PHF13,UGGT2,TSEN54,INAFM2,ZNF6
G1,CNN2,YWHAZ,ARHGDIB,PFN1,TMSB4XP4,ARPC1B,COTL1,MOB1A,RHOA,CAPZB,TLN1,C
NPO,TSPAN33,IER5,PEA15,SNX29,ANXA6,BCL2,PIEZO2,AUTS2,CYTH1,RAB29,HHAT,TNFSF4,
,IPCEF1,CHI3L2,HVCN1,PTPN12,ENTPD1,LBH,IRF8,PRKCH,DENND2D,ETHE1,MS4A7,NSMCE
C1LI2,PHKB,CSNK2A2,AMFR,PDPR,NLRC5,SLC7A6OS,MON1B,ATP6V0D1,ZFP90,GABARAPL
3,SOWAHC,FILIP1,ASPH,LINC02389,IRF4,EFNA5,ATP10D,AKAP6,ZNF462,GUCY1A1,NMT2
ASA1,ARID1B,SMAD3,TMEM38A,LIMD1,PRKX,HIVEP2,CCNL1,PSMA3-AS1,EZH1,TPCN1,ZNF
L1,EIF4EBP1,TRIB3,SHMT2,MTHFD2,AGA,SLC39A9,TTC39B,CTNNA2,SLC1A5,DDIT3,AL1578

CASTOR3,LINC00607,AC114495.2,AL133445.2,INPP5A,SMCO2,TMEM67,ALG14,STK32C,AL591

,IMPDH2,SLC25A3,ATP5MF,SOD1,NDUFB6,ACAT1,UQCRQ,HSD17B10,AKR1B1,ATP5F1A,NDU

XIST,VPS13A,ATG16L2,DIAPH2,NUP210,SEPT7P2,PABPC1L,AL450384.2,HERC2P2,NKTR,GOLG

4,NDE1,CD58,SNORD13E,NFKB2,CREB1,TGIF2,ZBTB21,CSF1,TIAM2,TRPM7,NFKBID,AC01193
1,SEC63,TMED2,TMED9,SELENOK,RPN2,TMEM50B,STT3A,TM9SF3,P4HB,MZB1,SURF4,LMA

3,SRSF1,PTGES3,HSPA4,SLIRP,EIF1AX,SET,PSMA3,MRPL3,AHSA1,EIF2S1,CDC123,PAICS,STI

3-AS1,AC131888.1,BCL11A,AC106870.1,RN7SL483P,LRRC74B,OR7C1,MTG2,THG1L,LUCAT1,D

F2C,ZBTB38,MBNL1,CDV3,CREB3L2,TCF4,TXNDC11,IGHJ3P,UBE2G1,SLC44A1,HS2ST1,VDR,
A80B,SNORD116-19,PLEKHH3,AC108704.2,RF00100,AC020916.1,LINC01814,PPP1R3E,AC01049
61A3,LINC02227,HLA-DQB2,LPXN,PARVG,ALDH5A1,SYNGR2,RGS13,COL9A2,ABCG1,HIBAD

NFRSF10B,FAM169A,AMZ2,HES2,FEZ1,ABCA12,APOBEC3C,INPP1,EDA2R,CCNK,DSE,CCNG1,
VDR76,POLD3,RFC2,TOPBP1,USP1,MSH6,DSCC1,GMNN,EZH2,GMPS,BRIP1,CHAF1B,SUPT16H

079466.1,APOL2,SLC2A3,ERO1A,ANKRD37,RLF,RBPJ,SLC2A1,DNM1L,AK4,WNT10A,PGM1,BN
AT,ARF1,FDPS,MRPL9,AHCTF1,FH,TIPRL,UCHL5,IVNS1ABP,HAX1,MTR,PRPF3,FLVCR1,SLC2
IF18B,KIF23,NUF2,CKAP2L,CIP2A,CKS1B,NCAPH,SHCBP1,BUB1,CCNF,KIF18A,RACGAP1,PR
IM1,UTRN,UBE2L6,STAT2,IGF1,PBX3,WASHC3,VNN2,TRIM38,STX7,ANTXR2,CMPK2,JAK1,L
Y,PSMD2,NDUFS6,UFD1,RBBP7,TIMM10,CENPW,HNRNPH3,MAGOH,MRPL51,LSM3,CALM1,S
A,RPL7A,AC131235.1,EEF2,RPLP2,RPS7,RACK1,RPS5,RPL12,RPLP1,RPS27A,RPL10,RPS8,RPL2
HP2,PRDX5,LAMTOR4,NADSYN1,FLOT2,NDUFV1,CIZ1,EXOSC5,TRPC4AP,PRPF6,COX5B,FB
.4,RF00019,ARHGEF37,CAPN10-DT,AL136295.5,TMEM116,ILF3-DT,LOH12CR2,AC133552.1,TC
IP1,POLR1C,UTP20,COA7,MAT2A,TCERG1,CTPS1,WDR12,DNAJC2,GNL2,RIOK1,RPL7L1,RSL
,NCF2,PASK,GBP4,KSR1,RAB9A,TNFAIP2,LINC00158,CCND2,BLVRA,CCDC28B,TNFRSF8,DD
S1,RAB3GAP2,NEK7,DUSP10,ADAR,RNF115,CDC73,YY1AP1,GATAD2B,GON4L,MIA3,ATF6,R

,KIAA0586,EMC9,WDR41,SRGAP2,ARHGAP27P1-BPTFP1-KPNA2P3,RBBP6,CCDC85C,CASP6,
3,VRK1,CENPP,FBXO5,MYBL2,CEP152,ASF1B,RAD51C,CCDC14,CENPO,EME1,HAUS8,TMEM
,RPL10AP6,RPLP0P6,RPSAP58,RPLP0P2,RPS15AP1,RPSAP54,AL136454.1,AC246787.1,RPL14P1,

NF26,SRGAP2C,KBTBD2,VANGL1,SCLT1,ZNF165,MYEF2,RALBP1,SUN2,WDPCP,MTIF3,YTH
TNRC6B,CYTIP,UBE2H,ZDBF2,CDKN1B,CHD2,CDC42SE2,AKAP13,BTG1,SEC14L1,MAN1A2,P

ORO1C,SKAP2,VASP,GBP1,RSU1,CLIC1,CFL1,IQGAP1,HDAC1,SNX3,CMTM6,CAPZA1,VIM,O
LYST,F11R,ITPKB,MACF1,HIPK2,IL6ST,ITPR1,NR3C1,RAB5B,KLHL6,TMEM131L,LRCH1,ETV
3,CREB5,SAMSN1,TLR10,SLC15A4,CDK5R1,BCL2L11,IL16,UST,NAALADL2-AS2,MAP2K6,AT
2,LONP2,WDR59,KLHL36,RSPRY1,CMTR2,CFDP1,COG4,ATXN1L,TCF25,BBS2,ADCY7,RANB

485.1,AC018462.1,MYRFL,AC040160.1,ZNF425,NR1H3,SLC12A8,CTBS,CC2D1A,AC083798.2,ZN

JFB3,ABRACL,PLS3,ATP5PO,ATP5ME,PRDX3,COX7B,ATP5MD,ATP5MPL,HNRNPC,CSTF3,OC

N1,OSTC,UGGT1,PDIA5,AC105250.1,FKBP2,SEC61A1,SRPRA,DERL2,FICD,SEL1L,MYDGF,TM

1,XRCC5,PSMA5,SSB,SRSF3,CYC1,PDCD5,FKBP4,PSMA7,RUVBL1,NUDC,PSME3,TIMM23,BC

DX25,CARD14,SRGAP3,Z95114.1,AC120042.1,PARVA,SHANK2-AS1,KLK2,FPR1,SLC35E1,AC00

OXO1,TUT7,PCM1,TRAM1,ARID5B,BTLA,CLEC2D,PABPC4,SLC1A4,CCDC88A,JMJD1C,DNAJ
.1,RNU2-63P,CASC19,TMEM107,LINC01225,DGKZ,SNORA65,AL117339.5,AC105105.1,RF00019,

CCDC90B,MIR34AHG,CDIP1,TP53I3,PHPT1,MACO1,ANKRD20A5P,SULF2,ZNF561-AS1,SESN1,
,ICMT,RFWD3,PRKDC,DUT,POLA1,SMC3,SUZ12,TCF19,CDT1,CENPU,MCM5,CHAF1A,TIPIN,E

IP3P1,AL354707.1,YEATS2,GOLGA8A,ARHGAP18,ZNF654,CERNA2,RNMT,AC097534.2,CCNG2
5A44,TARS2,POGK,DAP3,COA6,COX20,NUP133,PMVK,PYCR2,EEF1AKNMT,TRIM11,ALDH9A
1,SKA1,DBF4,SGO1,CEP55,TACC3,MAD2L1,SPC25,KIFC1,RAD21,KIF22,CKAP5,CDCA8,TMPO
PIN1,LINC01781,CCR1,SCPEP1,RB1,TMEM59,ITM2B,FCRLA,RNF213,ANKRD44,DDX60L,TRIB
TMN1,HNRNPL,KHDRBS1,LSM5,ANAPC15,TRMT112,PTP4A2,FKBP3,SNRNP200,ENY2,TUBB
,LRRC75A-AS1,RPS19,RPL19,RPL41,RPL22,RPL21,RPL5P34,RPL32,RPL29,RPL15,RPL36,EIF3F,
H1,HTT,TNIP1,CHID1,ZNHIT1,CINP,PI4KA,WDR46,SPTAN1,ACADVL,NF2,C12orf10,ROMO1,US
E3,AL355388.2,AC090114.2,RAB4B,AC091271.1,DNAJC3-DT,FKBP11,RALY-AS1,ZFYVE27,RF0
1D1,WDR3,RRS1,UMPS,MPHOSPH10,NOL6,PDCD11,GTPBP4,LTV1,POLR1B,AKAP1,WDR75,T
R2,FAM129A,FSD1L,VCAM1,PLEKHG1,NTRK2,DMXL2,OXTR,CYB5A,SNN,TTF2,TFDP2,IER3,
COR3,CD55,ACBD3,PRKAB2,SUCO,AL390728.6,RC3H1,SCYL3,ZBTB41,BROX,DCAF8,ARHGA

SPA17,DDAH2,FOPNL,KRT10,USP24,CRLF3,SAFB2,LNP1,KIF21B,SLCO3A1,CEBPZOS,EEF1AK
106C,AC007240.1,GPR19,C17orf53,C2orf48,MNS1,CEP57,GSTCD,CCDC15,LIN9,XRCC1,C5orf34,
RPS2P46,RPL21P16,AL590867.2,AL122020.1,RPL13AP25,RPS7P1,AC116533.1,RPL21P28,RPS7P3,

HF21A,PIK3IP1,RORA,UBR5,ATRX,SYF2,SERINC3,PINK1,EVI2B,MKRN1,UBR2,CCDC32,DNAJ

TF1,MSN,ILK,CAPZA2,PRKAR1A,GLIPR1,STK26,DCTN1,FIBP,PTBP3,HNRNPH2,NXT2,RAB7A
6,EPS15,BIRC2,IL13RA1,PIAS1,ORMDL3,SLCO5A1,ST3GAL6,PIK3R5,ATP6V1G1,ABCA5,CCSE
N1,SH3BP5,PHEX,PLXNC1,GDA,P2RY11,ZNF791,GPR82,CD52,CYFIP2,ARHGAP9,LCK,C12orf
10,ATMIN,SIAH1,ACSF3,PRMT7,ARL2BP,SPG7,MBTPS1,SNTB2,PLA2G15,GFOD2,MEAK7,ZN

)SF2,EMC7,SPCS2P4,UGDH,ARF4,NOMO1,PRRC1,SEC24C,ALG5,CNPY2,CDK2AP2,TMEM214,P

CIP,CCT8,VDAC1,SF3A3,NPM1,PHB,PA2G4,CMSS1,PRELID1,SNRPF,MRPL18,CHCHD3,GSPT1,

8764.3,MZT2A,ZNF549,ZNF114,CCDC88B,GPR1-AS,ANKS4B,KCTD20,RN7SL3,AC097376.2,AC0

B9,NBEA,SOS1,CCDC69,PEBP1,AC027290.2,S1PR1,TIPARP,FAM126A,MED13L,GPR15,IFNAR1,

ASTN2,ACTA2,TMEM68,CTNND1,TEX9,AC025423.1,CEP85L,PPM1D,ZNF79,KIAA1671,TMEM1
3LM,MMS22L,EXO1,GMPSP1,POLR3K,RFC3,DNA2,FANCA,TIMELESS,PRIM1,HAT1,RBL1,KNT

!,SNAPC3,PPME1,NOL3,PAM,CHSY1,MIR210HG,ANKZF1,MPI,PIK3R6,ZBTB25,WDR54,RNF24,A
.1,CSRP1,MRPS21,PRDX6,SRP9,SCCPDH,SOAT1,CHD1L,DARS2,CNIH4,ADSS,GPR89A,RBM8B,
),FAM83D,TTK,SPDL1,TICRR,ESPL1,CIT,NUSAP1,PLK4,DEPDC1B,BUB3,ERCC6L,POC1A,AC09
2,SHISA5,DTX3L,NT5C3A,RNF144A,PTPN22,SETBP1,TRIM25,LASP1,WDFY1,ARHGEF3,ANKRI
5,VPS29,UBB,CBX1,C12orf75,SEPHS1,GNAI3,PIN4,DNAJC8,ALYREF,GNB1,RBM42,CENPS,POLI
NACA,RPL37,QARS,RPL34,RPS13,SNHG5,EIF3H,RPL35A,EIF3E,BTF3,ERGIC3,IGBP1,UBA52,RF
SP22,CCT7,GNAI2,GSTP1,ACAA1,ARAF,HDLBP,PEX6,MTCH1,ACOT7,VPS51,ETFB,PNOC,SART
0019,AP000845.1,MAGIX,EIF2AK3-DT,AC107375.1,AC092119.2,AL645728.1,AC008764.7,AC0691
VNK,RIOX2,ESF1,DHX33,TWISTNB,TXLNG,USP36,NOL8,PDCD2L,CD3EAP,EIF3A,KNOP1,RBM
ABCC4,LMNA,NEDD4L,NBN,PLD1,TBC1D4,MBOAT2,EPS8,ARHGAP26,CDC42EP4,EPHX2,LAC
P30,RAB4A,CNST,POU2F1,POGZ,PRUNE1,UBE2Q1,SFT2D2,DSTYK,PPP2R5A,AIDA,GOLPH3L,(

,RPL10P9,RPL10P16,AC133134.1,AC079922.1,RPS27AP16,AC136632.1,AC000089.1,AC115223.1,R

B14,CIR1,PSEN1,SOS2,KLHL28,SLC12A6,YPEL2,RB1CC1,PELI1,PLAG1,ARID4A,SLU7,ARHGAI

R2,TSC22D3,SH3KBP1,LINC02384,SMAP2,ADAM22,WASHC2A,DHTKD1,MSC-AS1,CD84,CLNK
77,CD226,TWSG1,TMC8,HEG1,LINC01307,NCOA1,CAB39,PPFIBP1,UBE2F,ABI3,ENC1,NCR3,SL
F276,MARVELD3,CMIP,WWP2,CHTF8,TAF1C,TMEM170A,MLYCD,ZDHHC7,COG8,FTO,AC0091

NDUFAB1,HSP90AB3P,GLRX3,KPNA3,ABCF2,SNRPA1,HDAC2,CSE1L,HNRNPR,MTHFD1L,PS[illegible]

UBE2D2,FOXN3,TRAM2,ERN1,ETS1,LMO4,MANEA,CD47,ERP44,MAN1A1,PPP3CC,GPATCH11

68,NDUFAF6,BBC3,DRAM1,HSDL2,CSNK1G1,TIGAR,SMAD5,PLXNB2,PAPLN,F5,PRKAB1,DY[illegible]
[illegible]C1,POLE2,DONSON,BRCA2,GINS1,SRSF10,ORC6,MCMBP,BAZ1B,RIF1,GINS4,UHRF1,CENPK,[illegible]

[illegible]L133453.1,MXI1,VGLL4,SDAD1P1,AP000769.1,RAB42,KDM4B,DARS-AS1,NR2F2-AS1,FBXO42
INTS7,RBBP5,ADIPOR1,UTP25,TSEN15,PPP1R15B,UBQLN4,SMYD2,PRCC,COG2,JTB,C1orf131,[illegible]
[illegible]1057.1,CDCA3,CCDC18,MZT1,CDK5RAP2,PARPBP,NDC1,FBXO43,MIS18BP1,CENPI,SUV39H1,[illegible]
[illegible]13A,WIPF1,DHX58,ISG15,RCBTB2,HELZ,EVI2A,BISPR,ARHGAP15,DCAF5,AL096799.1,DMXL

[illegible]L22P1,RPS7P10,SNHG6,RPS4XP6,RPS4XP11,RPL12P4,RPS26,AL139095.2,RPS3P6,SLC25A6,RPL
[illegible]1,PPP1R7,ARID3B,UXT,REC8,ELOF1,NAA10,INPP5K,SMARCD2,TMEM248,PYGB,TECRP1,TA[illegible]
85.1,WDR5B,LINC01560,AC012360.3,AC023157.3,AC017083.1,PAXIP1-AS1,GABPB1-IT1,AC2398
[illegible]125,AMD1,PPRC1,SNHG15,SNHG17,RRP9,DNTTIP2,PPAT,NAT10,WDR36,SLC39A14,RRP15,PR[illegible]
[illegible]TB,GPHN,ANXA7,IGSF3,DMD,HEPH,AHCYL2,TUBB2A,PRAME,RHOV,MREG,GRIN2A,TNFRS[illegible]
[illegible]LK2,KCNK1,SLAMF6,LPGAT1,BLZF1,MEF2D,VAMP4,ODR4,SSR2,PIGC,BCL9,AC096642.1,SE[illegible]

PL10P3,RPL37P2,AC034236.1,AC024293.1,RPS11P5,RPSAP61,RPL4P5,RPL26,RPL7AP6,AP00102[illegible]

,TIFA,WHRN,EZR,PITPNC1,ABCA10,PACS1,NEXMIF,CDK17,ABCC1,ZC3H12D,PPARA,LRCH3,

[illegible]20.2,AC007342.6,AC079416.1,ZNF778,SLC38A7,ZNF319,C16orf70,DHX38,ZFP1,ZNF19,CENPBD1

MD11,CHORDC1,DDX1,IARS,LAS1L,TFAM,NUP93,MRPL21,PSMD12,DCAF13,PSMD6,GTF3C6,[illegible]

,HOOK3,SPTY2D1,ERLEC1,WIPI1,CUL3,NT5C2,ST6GALNAC4,PEX2,TMEM19,LYSMD3,COG3,[illegible]

NC1H1,CAV2,RAB38,PGAP1,TP53TG1,RETSAT,F2R,ITM2A,BAX,SORD2P,RINL,RAB39B,TMTC
CDCA7L,CDK4,CTNNAL1,RMI1,ORC1,GGCT,KLHL23,EXOSC9,RFC1,FIGNL1,FKBP5,CDC7,DSN

,AC023632.6,PDK3,DUSP3,MYLIP,MLLT3,MINDY2,ARL10,SIT1,TMEM263,TBC1D22A,PLOD1,S
TOR3A,MPC2,DUSP12,UAP1,MPZL1,KCTD3,CHTOP,TSNAX,CREG1,RABIF,UCK2,POLR3C,TFE
SAP30,CCDC34,CENPJ,CCDC150,C1orf112,SKA2,SPC24,CTCF,TRIP13,TRAIP,PKP4,CNTRL,CEN

41P2,RPS20P14,CCNB1IP1,RPS28,RPL18,RPL5P1,RPS17P5,RPL7P23,AC098934.3,CSDE1,TPT1P9
DA3,CCNY,TECR,ATPAF2,CUL9,NDUFA11,CCDC107,BIN3,PHF23,IP6K1,AMPD2,KMT2B,DRG2,
68.1,FNBP1P1,HIST2H2BD,AC098591.2,AL358472.5,TBCE,ERV3-1,MIAT,SIRT7,SCAMP4,AL121
F38B,URB2,UTP15,GEMIN5,URB1,UBE2G2,MIEF1,DDX10,NAA25,QTRT2,LARP4,BYSL,UTP14
F9,ANKLE2,NID1,RDX,RSPH1,CD9,NCK2,HEY1,FBN1,NOL4L,STAT5A,EHBP1,GNG2,CSF2RB,N
TDB1,XPR1,NCSTN,ZBTB18,ANGEL2,PI4KB,ARNT,BX571818.1,ZBTB37,GPATCH2,ARL8A,DEN

.1,RPL26P6,RPS19P3,AC069213.3,AC004453.1,NACA3P,RPS3AP49,NACA4P,AC090114.1,AL4504

LRRC59,YWHAE,GTF3A,SNRPB2,FAM136A,STOML2,HSP90AB2P,PSMC2,RARS,POLR2F,AK2,T

N1,NUP155,DCLRE1B,RPA1,DMC1,POLE,CDK2,AL161891.1,MASTL,SVIP,PAXIP1,NUP85,WEE1

32M,SPRTN,TAF5L,AC009487.2,SCNM1,DESI2,TADA1,FAM20B,YOD1,SCAMP3,GUK1,NVL,CEI
PL,RHNO1,RCCD1,GEN1,CEP128,POLH,TDP1,H2AFX,NIF3L1,C2orf69,GMEB1,OIP5,Z94721.3,NI

,EEF1A1P11,AC069218.1,EEF1A1P5,AL050331.1,RPS4XP7,EEF1A1P6,RPS4XP13,AC084824.1,EEI
,EXOC3,FAM168B,CHKA,NPLOC4,TMEM8A,TMEM222,PRKD2,DUS2,NMRAL1,PGS1,METTL26
558.1,SFT2D3,SWSAP1,EIF3J-DT,PGBD4,THUMPD3-AS1,PIGV,KRT8P12,AC144652.1,AC002310.
A,PTRH2,PUS7,POLR3E,BAZ1A,MAK16,TSEN2,PNO1,TRIM27,DPH2,ZNF121,ALKBH2,RRP1B,F
NEURL3,LYSMD2,POU2F3,AL133330.1,TNFRSF11B,KALRN,HNRNPA1P21,TMEM170B,ACKR3,V
NND4B,RGL1,TBCE,ZNF496,GPR89B,GALNT2,C1orf56,RRNAD1,NME7,SLC41A1,CEP170P1,CYB

405.1,RPS3AP25,RPS13P2,AC016739.1,RPL31P12,CCDC88C,RPL5P4,RPL13P12,RPL15P3,RPL23AI

TFRC,MRPL15,MTCH2,HSPD1P1,GTF2A2,CAPRIN1,IDH3A,RWDD1,NDUFAF2,OLA1,LETM1,VD

,BARD1,RECQL,RAD18,PARP2,PRIM2,RPA2,CDCA4,SLFN13,TEX30,CASP2,COPS3,NUP153,KD

RS2,ARV1,GLRX2,COX20P1,FLAD1,B3GALNT2,TAF1A,THEM4,CHML,ZNF670,HNRNPUP1,C1o
EIL3,MPHOSPH9,KATNA1,RF00019,INCENP,PHF19,TBC1D31,CHEK2,CCDC77,TRIM59,ARHGE

F1A1P13,RPL26P19,RPS4XP16,SEC11A,HNRNPA1L2,RPL41P1,LETMD1,RPL23P8,EIF3FP3,RPL7
,CHST10,KLHDC3,PAFAH1B3,GPR108,IPO4,AC079250.1,UPF1,PPP4C,MRPL2,AP3D1,MAD2L2

ASTKD2,POLR1A,UBIAD1,IMP3,DDX31,SRFBP1,GTF3C4,SURF6,MPP6,RCL1,PLD6,XPO5,C12or
VLDLR,LHX2,IL17RB,TTYH2,TMEM178B,AL390755.1,CETP,CASK,CFP,AC008691.1,DPYSL2,GC

P42,RPL24P8,RPS21P4,AL355032.1,RPSAP19,AC099789.1,AL033519.2,AC008850.1,RPS2P35,RPSA

AC2,EFTUD2,PPIL1,RNASEH1,KRR1,RPL22L1,USP14,PPID,MRPS23,PDAP1,PHB2,SEH1L,MRP

ELC2,CENPQ,BDH1,L2HGDH,NR2C2AP,RBM17,PMS1,HMGXB4,LRRC20,RFC5,SKP2,HAUS6,E

F39,IQGAP3,TAF5,CEP44,C18orf54,AC091057.4,CENPC,HASPIN,FAF1,MTFR2,CDKN2C,RSRC1,

P32,AC113367.1,RPS4XP2,AL049873.1,RPS3AP5,RPL5P17,RPS2P5,RPS15AP11,RPS15AP38,AC11

f29,PMM2,ZNHIT6,UBTF,MTPAP,NAF1,ZC3H8,ZNF485,RPL36A,SDAD1,URI1,METTL2B,TNPO
NT1,SERPINB10,LNX1,PRRX1,B4GALT6,BATF3,CRYBG1,STMN4,FLNB,MED14,DAPK1,MTE

P18,AL590682.1,RPL10P6,RPS18P9,AC092128.1,RPS2P7,RPL13AP20,AC004057.1,RPL23AP65,RP

L14,MRPL35,HCCS,LLPH,METAP2,PSMC1,ELOC,NABP2,EIF2B3,POLR2K,PSMC5,PSMC4,CCDC

MC3-AS1,LUC7L2,RAD1,AC022182.3,FANCB,RCC2,UBA2,PHF10,LIG1,TUBGCP3,EXO5,CAND1,

ASF1A,CWF19L2,CETN3,DBF4P1,C12orf65,DLEU2,CDR2,ING1,RDM1,AC015849.5,PHTF2,KMT5

0994.2,RPS9,RPL34P18,TPT1P4,EPB41L4A-AS1,RPL29P11,ZNF581,ZNF277,RPL5P32,AC091429.1,

2,TGFBRAP1,DDX51,DDI2,TRMT10C,ADAT2,DCAF1,KLHL8,PER2,WDR74,FP565260.1,POLR3D
RF4,HUWE1,TIMP1,SNX7,MERTK,PAQR8,ARNTL2,NHSL1,SLC6A4,LAMB3,AFAP1L2,IL2RA,SP

L12P38,RPSAP9,RPSAP3,RPS18P12,RPL7P6,AC020898.1,RPL5P12,AC097658.1,AC091042.1,EEF1

:58,NSFL1C,SMARCA5,METTL5,LRPPRC,RPL26L1,TXNL4A,BCAS2,G3BP2,TKT,TRAPPC4,MRI

,JOSD1,RNF219,MLH1,SHMT1,SLF1,SUPT16HP1,BTG3,FANCC,UBAC1,HOOK1,TONSL,GINS3,N

A,XRCC4,ZNF850,HNRNPUL1,MED30,PKNOX1,UBL7-AS1,HIRIP3,SRBD1,MED21,COIL,TMPO-

,CEP83,PRMT3,NKRF,SUPV3L1,INO80,ZNF146,AC015802.6,TYW1,BOD1,PUS1,UBP1,SLC19A1,S
ART,PTPN1,MMD,PIWIL4,AL390719.1,PRLR,CYB5R2,ANXA3,PARM1,IL3RA,RALA,FSCN1,CLE

GP5,AC068631.3,RPL21P116,RPL7AP50,AL022718.1,AC107956.1,RPS2P55,RPL9P32,RPL36AP39,

PL19,DLAT,MRPS18C,MRPL16,MRPL20,MRPL47,DFFA,DYNC1LI1,ACLY,AIMP2,UQCRFS1,SAM

NUP160,NUP58,PM20D2,CHRNA5,TOP3A,TUBGCP5,SRSF4,SFMBT1,C4orf46,SFXN1,NPAT,ADC

SNORA74D,GTF2H2,ZNF202,METTL16,PUS3,METTL1,NMD3,DGAT2,MARS2,BAG1,TIMM9,UT
C17A,CLDN16,JPH4,FLVCR2,FHOD3,GRHPR,OSBPL3,LMO3,RAP2A,HMSD,CELSR1,MARVEL

[illegible]2,LHFPL6,PLAU,NPAS3,MTMR4,KYNU,L3MBTL4,HMGN3,TPCN2,ITGBL1
