## Supplementary material for "Human transcription factor combinations mapped by footprinting with deaminase": Table S2

| | DddB (BadTF3) toxin-Immunity protein complex |
|---|---|
| **Data collection** | |
| Wavelength (Å) | 1.541838 |
| Space group | C2221 |
| Cell dimensions | |
| a,b,c | 84.1, 130.6, 61.6 |
| α,β,γ | 90, 90, 90 |
| Resolution (Å) | 12.62-2.50 (2.59-2.50) |
| Rmerge (%) | 7.5 (29.8) |
| I/σ | 25.4 (7.8) |
| Completeness (%) | 99.94 (99.58) |
| Total No.of reflections | 116898 (11407) |
| Unique reflections | 11988 (1175) |
| Redundancy | 9.8 (9.71) |
| **Refinement** | |
| Resolution (Å) | 12.62-2.50 |
| No. of reflections | 11971 |
| Rwork/Rfree (%) | 19.33/23.76 |
| No. of atoms | |
| Protein | 1780 |
| Ligand/ion | 1 |
| Water | 95 |
| Average B-factors (Å$^2$) | 37.04 |
| rms deviations | |
| Bond lengths (Å) | 0.009 |
| Bond angles (°) | 1.03 |
| Ramachandran plot | |
| Favored (%) | 98.18 |
| Allowed (%) | 1.82 |
| Disallowed (%) | 0 |
